# Hyperextended telomeres promote C-circle formation in telomerase positive human cells

**DOI:** 10.1101/2023.01.26.525615

**Authors:** Celina Y. Jones, Christopher L. Williams, Sara P. Moreno, Danna K. Morris, Chiara Mondello, Jan Karlseder, Alison A. Bertuch

## Abstract

Telomere length maintenance is crucial to cancer cell immortality. Up to 15% of cancers utilize a telomerase-independent, recombination-based mechanism termed alternative lengthening of telomeres (ALT). The primary ALT biomarker is the C-circle, a type of circular DNA with extrachromosomal telomere repeats (cECTRs). How C-circles form is not well characterized. To investigate C-circle formation in telomerase+ cells, we studied the human cen3tel cell line, in which telomeres progressively hyper-elongated post *TERT*-immortalization. cECTR signal was observed in 2D gels and C-circle assays but not t-circle assays, which also detect cECTRs. Telomerase activity and C-circle signal were not separable in the analysis of clonal populations, consistent with C-circle production occurring within telomerase+ cells. Two other long telomere, telomerase+ (LTT+) cell lines, HeLa1.3 (~23 kb telomeres) and HeLaE1 (~50 kb telomeres), had similar cECTR properties. Telomerase activity did not directly impact C-circle signal in LTT+ cells; instead, C-circle signal correlated with telomere length. LTT+ lines were less sensitive to hydroxyurea than an ALT+ cell line, suggesting that ALT status is a stronger contributor to replication stress levels than telomere length. Additionally, FANCM did not suppress C-circles in LTT+ cells as it does in ALT+ cells. Thus, C-circle formation may be driven by telomere length, independently of telomerase and replication stress, highlighting limitations of C-circles as a stand-alone ALT biomarker.

## INTRODUCTION

With rare exceptions, cancer cells employ a mechanism of telomere length homeostasis to enable continuous proliferation over time (1). Approximately 85-90% cancers activate or upregulate telomerase, an enzyme which is suppressed or very minimally expressed in normal somatic cells, to achieve telomere maintenance (2). A smaller proportion of cancers (~10%) utilize ALT pathway, which is telomerase-independent and instead involves a break-induced replication-like mechanism (3–5). The prevailing model describes telomerase-mediated extension and ALT as mutually exclusive within primary cancer cells. Determining which telomere maintenance mechanism (TMM) is present, if any, is clinically relevant as they can associate with different prognoses in various cancer types and are potential therapeutic targets (6,7).

Aside from telomerase activity, various properties differentiate the two TMMs. ALT+ cells are characterized by very long bulk telomeres, with individual chromosome ends displaying significant heterogeneity, elevated levels of telomere sister chromatid exchanges, and enrichment of telomeric sequence varied from the canonical TTAGGG repeat (3,8–10). They also have the feature of promyelocytic leukemia (PML) nuclear bodies associated with telomeric DNA, referred to as ALT-associated PML bodies (APBs), which are required for ALT-mediated telomere maintenance (11,12).

ALT+ cells also specifically harbor circular forms of cECTRs (13,14). Telomeric (t)-circles are a class of cECTRs that are most often characterized as nicked or gapped double-stranded circles. T-circles are typically detected by neutral-neutral 2-dimensional gel (2DG) electrophoresis where they migrate as a distinct arc (13–15), although this arc may also include looped telomeric sequence fragments (16). As a more sensitive method, the t-circle assay (TCA) was developed (17). Phi29 polymerase, which contains strong strand displacement activity, is used to amplify cECTRs after denaturation and annealing of a strand-specific primer. The product is a band that runs above bulk telomeric DNA following 1-dimensional gel electrophoresis. Due to its dependence on rolling circle amplification and on the denaturation and primer annealing steps, the TCA only requires that the cECTRs have a fully intact circular strand.

A distinct form of cECTR is the C-circle, which is defined by an intact CCCTAA telomeric repeat-strand that is partially double-stranded. C-circles are considered a biomarker of ALT, with the C-circle assay (CCA) serving as a robust and quantitative assay to identify ALT+ cell lines and tumors (18,19). Like the TCA, the CCA depends on phi29 polymerase amplification, but the maintenance of native conditions, absence of primer addition, and sequence specificity of the probe limit amplification and detection to intact circles with telomeric C-strand repeats that are self-primed (partially double-stranded). When the corresponding primer and probe are employed, TCAs should similarly identify the structure detected by CCAs.

The factors that regulate t-circle and C-circle production have been of subject to investigation. Several studies support T-circles arising from the trimming of extended telomeres. For example, t-circles are induced in telomerase+ cancer cell lines in association with telomere elongation driven by telomerase upregulation (15). They are also present in mature human sperm with elongated telomeres and mitogen-stimulated T-lymphocytes in which telomerase is upregulated and telomeres lengthen (20). Several factors have been shown to regulate t-circle elaboration in telomerase+ cells with extended telomeres or ALT+ cells, including Nbs1, XRCC3, ORC2, and TZAP (20–24).

Factors that have been identified to govern C-circle production to date have been associated with telomere replication stress. For example, SLX4 and FANCM suppress while BLM and other members of the BLM/TOP3A/RMI complex promote C-circle formation in concert with their effects on ALT-telomere synthesis (12,25–27). MRE11 and RAD51 also suppress C-circle formation in ALT+ cells in a RAD52-independent telomere synthesis pathway (28). In telomerase+ lines, reduction of the replication fork remodeling enzyme SMARCAL1 or the histone chaperone ASF1, which induces replication stress, promotes C-circle formation (29,30).

Tying the above concepts together, both t-circles and C-circles have been detected in human embryonic stem cells (hESCs) with t-circles generated by a Nbs1/Xrcc3-regulated trimming of elongated telomeres and C-circles promoted by increased replication stress, such as by treatment of cells with hydroxyurea (HU) (24).

Here, we have exploited the unique cen3tel human cell line to study cECTR production in telomerase+ cells. The cen3tel line was derived from primary skin fibroblasts obtained from a female centenarian immortalized with *TERT*-containing retrovirus (31). The cells were cultured long-term to examine cellular changes that accumulate post-immortalization. Following an initial period of telomere shortening through population doublings (PDs) ~80 – 120, telomeres continuously elongated to ~100 kb at PD1000, thus exceeding over 10-fold the more typical telomere length of a wide range of human cells. Telomerase activity similarly increased across PDs; though, it plateaued around PD600. APBs were absent; therefore, it was concluded that telomere lengthening was solely due to telomerase-mediated extension (31). During propagation, cen3tel cells were viably frozen at regular time points, establishing isolates with a range of telomere lengths (31). We analyzed these cells for the impact of excessive telomere length on cECTR production and found cECTR signal in cells with hyper-elongated telomeres in 2DGs and CCAs but, surprisingly, not TCAs. Similar results were obtained for two other LTT+ cell lines. To understand the contribution of telomerase to C-circle production, we knocked out and then reintroduced *TERC*, an essential integral component of telomerase, and found C-circle production was not influenced by telomerase directly but rather correlated with its impact on telomere length. Finally, we assessed the effect of HU-induced replication stress on cell growth and observed the LTT+ cells were not as severely impacted as ALT+ cells. FANCM depletion did not affect C-circle levels in the telomerase+ cells, indicating that different mechanisms underlie their production in telomerase+ versus ALT+ cells. Together, this work firmly extends the context of C-circles from solely ALT+ cells to somatic cells with telomeres that have been hyperextended by telomerase and demonstrates differences in potential contributors to their formation.

## RESULTS

### The continuous extension of telomeres in cen3tel cells is distinguishable from the heterogeneous lengths of ALT+ cells and the stable lengths of *TERT*-immortalized BJ (BJtert) fibroblasts

To further characterize the telomere extension phenotype of cen3tel cells, we compared their length distributions to those of ALT+ U2OS cells by TRF analysis (Figure 1A). Cen3tel cells at later PDs (420 and higher) had extremely long telomeres, with bulk averages ranging from ~30 – 100 kb. Like U2OS telomeres, some telomeric signal was trapped in the wells, likely tangled fragments of extremely long telomes. In contrast to U2OS telomeres, which had similarly very long (average ~50 kb), the Cen3tel cells lacked detectable short telomeres by TRF analysis, distinguishing them from classic ALT+ TRF profiles.

**Figure 1.**
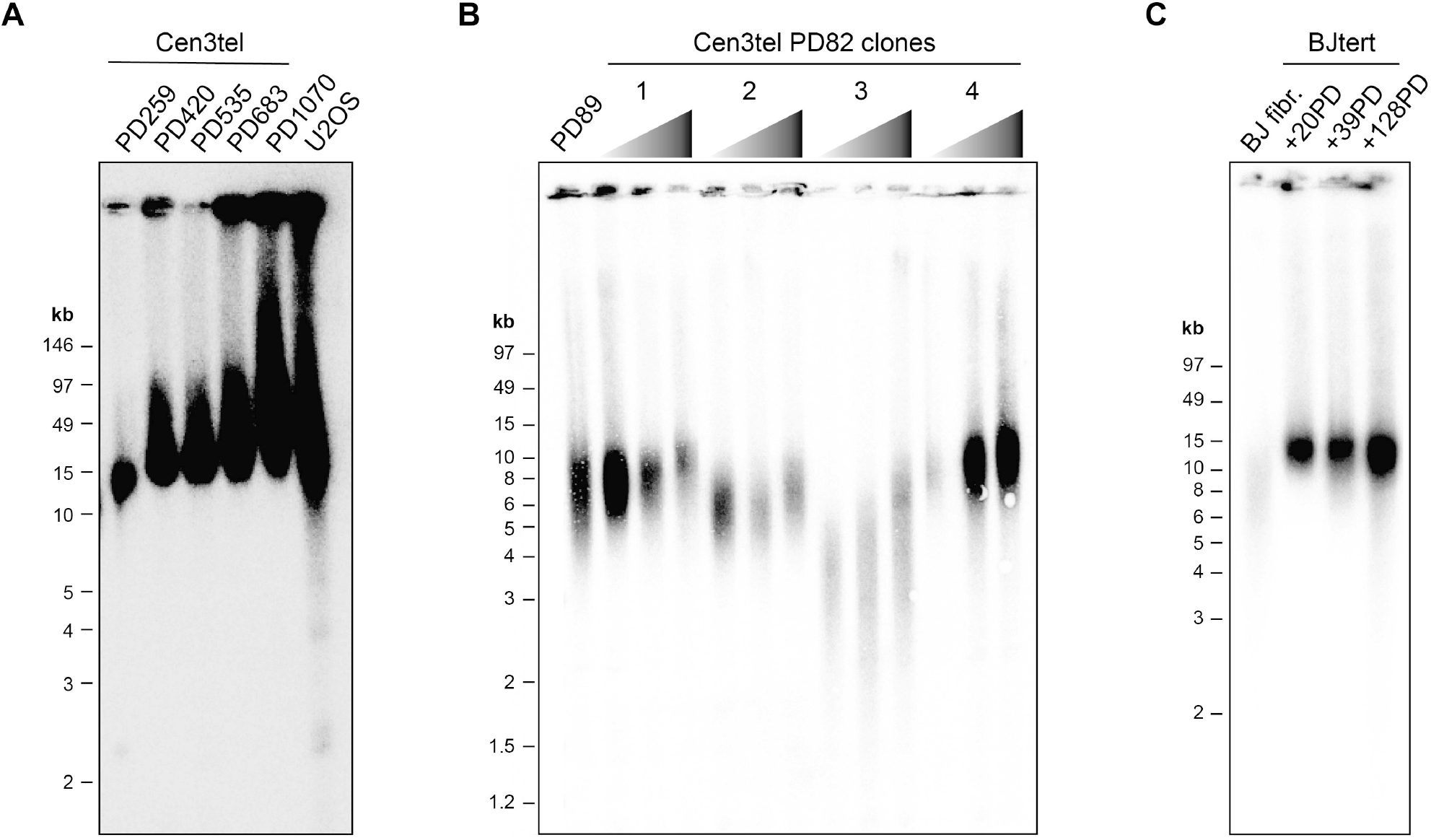
The continuous extension of telomeres in cen3tel cells is distinguishable from the heterogeneous lengths of ALT+ cells and stable lengths of *TERT*-immortalized BJ (BJtert) fibroblasts. (A) Telomere restriction fragment (TRF) assay using cells embedded in agarose plugs to maintain DNA integrity and pulsed-field electrophoresis (using PFGE certified agarose) to better separate very long TRFs. TRFs of the designated mid to late population doubling (PD) cen3tel cells are compared to those of ALT+ U2OS cells. (B) TRFs of clonal populations isolated from cen3tel PD82 cells grown an additional ~35, ~65, and ~95 PDs compared to those of a polyclonal sample at a similar PD. (C) TRFs of non-immortalized BJ fibroblast (fibr.) and BJtert cells collected at designated PDs following *TERT*-immortalization. See also Figure S1.

We next assessed whether the progressive telomere lengthening that occurred upon development of the cen3tel line would consistently replicate with individual cen3tel clones isolated at an early PD or similarly occur in other *TERT*-immortalized fibroblast cells. First, clonal populations of cen3tel cells were isolated from cells at PD82, a point before the line began telomere extension (31), and cultured an additional ~100 PDs. The starting telomere length in each clone varied, but telomeres extended over this period in all populations (Figure 1B). In contrast, while the telomeres in *TERT*-immortalized BJ fibroblast (BJtert) cells were longer than the non-immortalized BJ cells, they remained constant in length over a similar period of propagation as the cen3tel clones (Figure 1C). The lack of continuous telomere elongation in BJtert cells was not due to insufficient telomerase activity as it was comparable to that of the PD82 clones and did not increase substantially over the time period tested (Figure S1). This contrast highlights that cen3tel cells possess the capacity for continuous telomere extension after exogenous telomerase expression, superseding typical telomere length homeostasis mechanisms, and that this extension does not always occur in TERT-immortalized fibroblasts.

### Some but not all cECTRs characteristic of ALT+ cells are detectable in cen3tel cells

Although cen3tel cells lack APBs (31), their extremely long telomeres led us to determine if they had cECTRs. We employed the three commonly used assays, 2DG electrophoresis, TCAs, and CCAs. Starting with cells at PD181, we observed a cECTR signal in 2DGs where t-circles migrate, with the signal increasing with telomere length (Figure 2A), similar to the findings with HT1080 cells transduced to overexpress telomerase (15). In addition, there was an arc below the cECTR arc, where supercoiled telomeric DNA would be expected (32), in PDs 708 and 1104. Strikingly, despite the prominence of these arcs, the cen3tel cells lacked signals in TCAs (Figure 2B), regardless of whether the G- or C-rich strand was amplified (Figures S2A and S2B). These results demonstrate that, while there is extensive telomere elongation in the later PD cen3tels, they do not result in t-circles as detected by TCAs and, underscore that, despite 2DGs and TCAs often being used interchangeably to assay for t-circles, the structures identified by these methods can be distinct.

**Figure 2.**
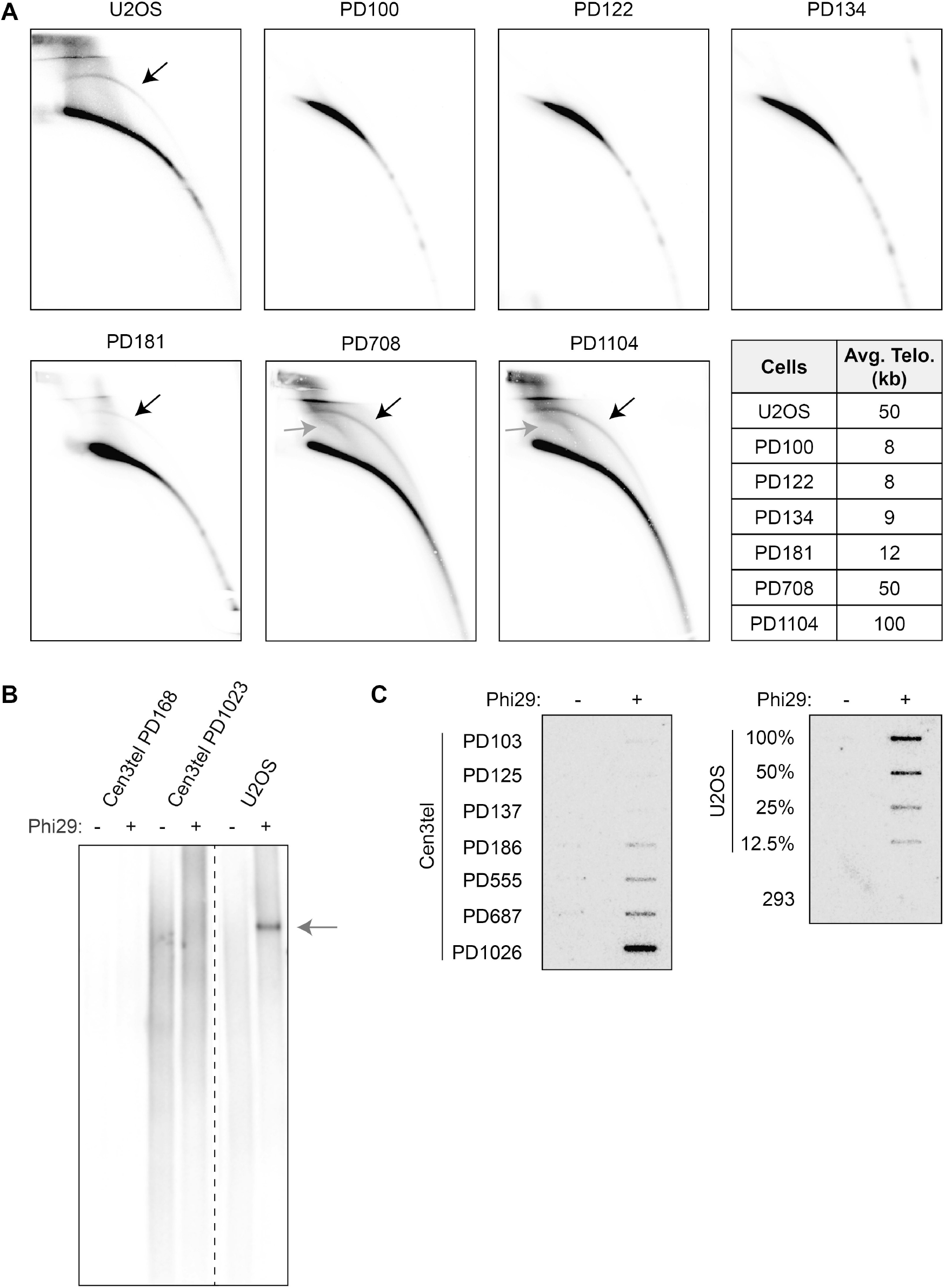
Circular extrachromosomal telomeric repeats (cECTRs) are detected in later PD cen3tel cells in some, but not all, cECTR assays. (A) Analysis of cECTRs in U2OS and cen3tel cells at designated PDs by 2D gel electrophoresis using a radiolabeled C-rich telomere probe. Black arrows note cECTR arcs; dark gray arrows note supercoiled telomeric DNA. Average bulk telomere lengths for each cell line are shown. (B) Analysis of cECTRs via t-circle assays (TCAs) using a C-rich primer and a digoxigenin-labelled G-rich telomere probe to detect the product. Arrow notes cECTR amplified product. (C) Analysis of cECTRs via C-circle assays (CCAs). Left panel: products of cen3tel cells from designated PDs. Right panel: products of serially diluted U2OS cells and 293T cells, which serve as positive and negative controls for C-circles, respectively. See also Figure S2.

In contrast, the cen3tels with elongated telomeres did produce a signal in CCAs, indicating the presence of C-circles (Figure 2C). The C-circle signal was evident at the same PDs as when cECTRs were found in 2DGs. The C-circle signal increased with telomere length. Moreover, at the latest PD, the intensity of the signal was similar to that of U2OS cells, which was different from prior reports of C-circle signal in hESCs or TERT-immortalized IMR90 cells, which were far less than those of U2OS cells (24,33). Retention of the C-circle signal after exonuclease digestion of linear DNA revealed that the templates for amplification were indeed circular (Figure S2C). These results underscore that t-circles and C-circles are produced by different mechanisms and C-circles can be produced at high levels in differentiated somatic cells with hyperelongated telomeres.

### Telomerase activity and C-circle signal occur simultaneously

The unexpected finding of prominent C-circle signal in cen3tel cells with robust telomerase activity could reflect each arising from separate populations of cells within the polyclonal cultures. To assess this, we isolated clonal populations at various PDs: before telomere elongation began (PD82 and PD92), shortly after elongation began (PD166), and after elongation was extensive (PD626 and PD1012). If our initial observations were due to a mixed population, we would expect that some, if not all, clones would display only telomerase activity or C-circle signal, not both. However, we found all clonal populations displayed both telomerase activity and C-circle signal (Figure 3). The telomerase activity level was qualitatively comparable to that of polyclonal populations (Figure 3A), as was the C-circle signal, whether the polyclonal population lacked or had robust signal (Figure 3B). TRF analysis showed that the clones’ telomere lengths were variable but generally within the range of lengths in the polyclonal populations (Figure S3). These data reveal that telomerase activity and C-circle production were not separable, suggesting the co-occurrence was not due to heterogeneous cellular subpopulations.

**Figure 3.**
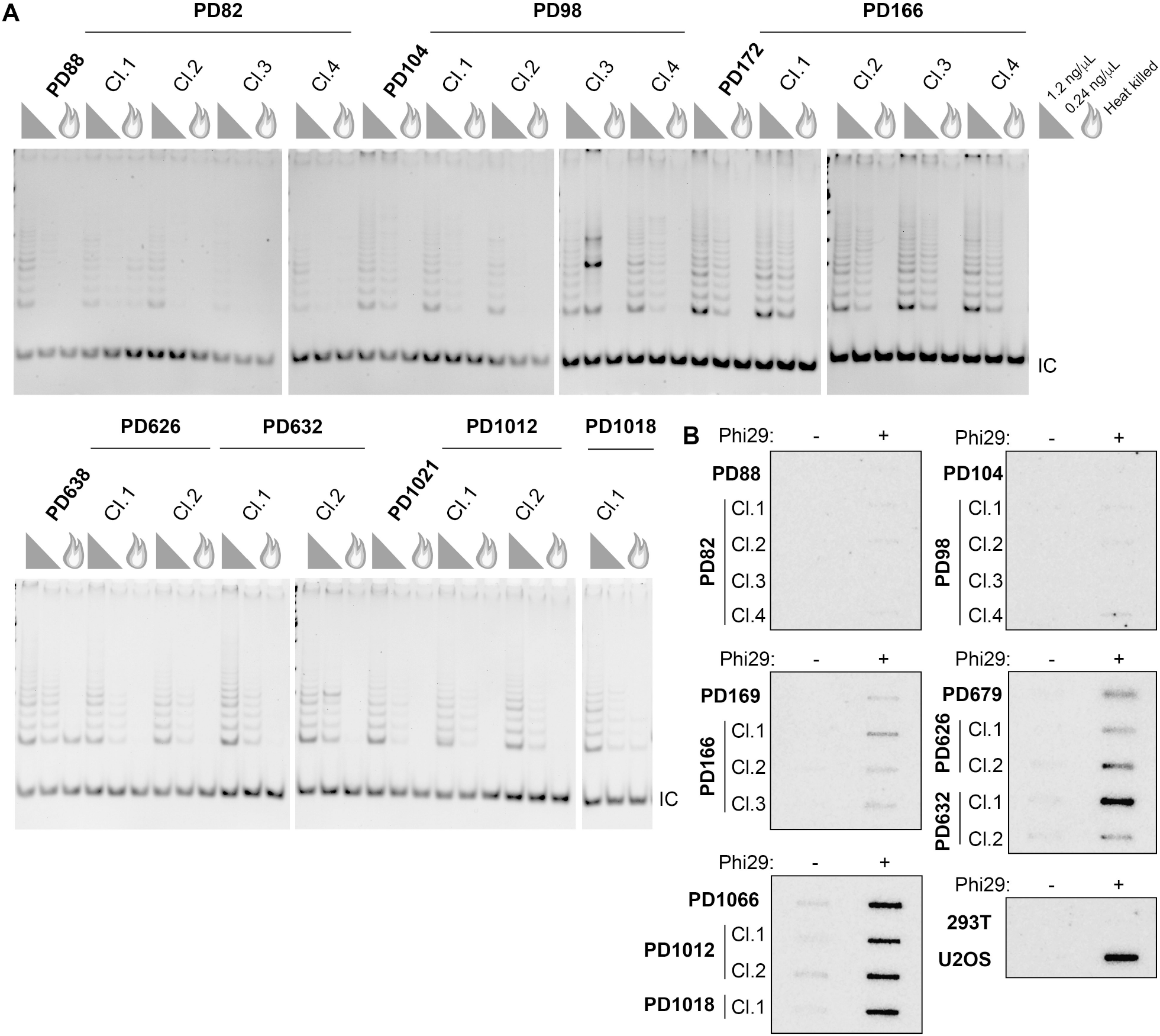
Both telomerase activity and C-circle signal are detected in later PD cen3tel clones. (A) Telomeric repeat amplification protocol (TRAP) assays of polyclonal populations and clonal populations isolated from the designated PDs. Assays performed with 1.2 ng/μl or 0.24 ng/μl protein concentration or heat-killed 1.2 ng/μl lysate (flame symbol). IC, internal amplification control. (B) CCAs on the same polyclonal and clonal cells as in A, with 293T and U2OS, negative and positive controls, respectively. See also Figure S3.

### cECTR generation in other telomerase+ human cells with long telomeres is similar to that of cen3tel cells with long telomeres

Given that cECTRs were detectable in 2DGs and CCAs but not TCAs in cen3tel cells with long telomeres, we assessed whether other telomerase+ cell lines with long telomeres had similar properties. Two HeLa cell lines are used widely as long telomere models. HeLa1.3 cells have bulk telomere lengths of 23 kb and were established by successive rounds of subcloning of the long telomere HeLa line, HeLa-L (34). HeLaE1 cells, on the other hand, were established by overexpressing both hTR and hTERT, creating a super-telomerase cell line with telomere lengths averaging 50 kb (35). To compare these lines with cen3tel cells of fairly comparable telomere lengths, we paired HeLa1.3 cells with cen3tel ~PD270, and HeLaE1 with cen3tel ~PD640 (Figure 4A). HeLa1.3 and HeLaE1 cells had robust telomerase activity similar to that in cen3tel cells PDs 172 and later (Figure 4B).

**Figure 4.**
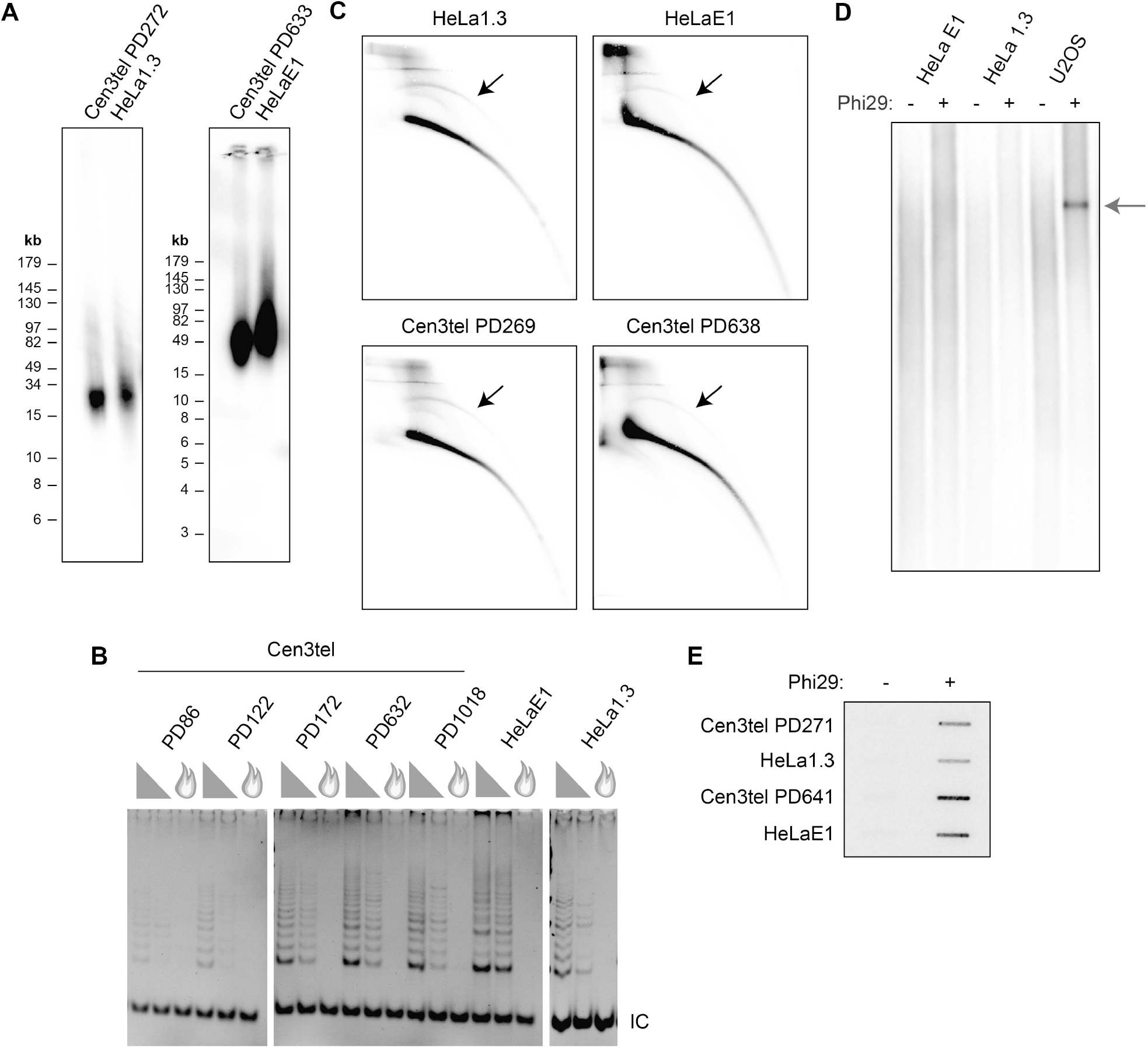
cECTRs are detected in long telomere telomerase+ (LTT+) cells in assays similar to those of cen3tel cells. (A) TRF analysis as in Figure 1 of the indicated cell lines of comparable telomere lengths. (B) TRAP assays of the designated cell lines. Protein concentrations used in assays as in Figure 3A. (C) 2D gel analysis as in Figure 2A. Arrows denote cECTR arcs. (D) cECTR analysis by TCAs as in Figure 2B; arrow marks the amplified cECTR band. (E) CCAs as in Figure 2C comparing cell lines of comparable telomere lengths. See also Figure S4.

We first examined cECTRs by 2DG electrophoresis and observed a cECTR arc in the HeLa1.3 and HeLaE1 samples at signal intensities similar to the corresponding cen3tel PDs (Figure 4C). As with the cen3tel cells, we did not detect cECTR signal by TCA (Figure 4D). Finally, both HeLa1.3 and HeLaE1 contained C-circles, with signals similar to the corresponding cen3tel PDs (Figure 4E). The C-circle signals in the HeLa lines were also resistant to exonuclease digestion (Figure S4). Thus, these assessments revealed that two other telomerase+ cell lines with long telomeres have cECTR characteristics similar to those of the cen3tel cells with long telomeres, suggesting that they were not exclusive to cen3tel cells. It is noteworthy that C-circles were detected in each of these telomerase+ lines, as the HeLa lines are cancer cell lines, and the cen3tel cells are immortalized primary cells, though the cen3tel cells adopted tumorigenic properties across propagations (31). Given their similarities, we grouped these long telomere, telomerase + lines together with the term LTT+ cells and will refer to them as such moving forward.

### Telomerase activity does not impact C-circle production in LTT+ cells

While considering what might underlie the elaboration of C-circles in the LTT+ cells, we focused on two qualities that distinguish these cells from typical telomerase+ cells: strength of telomerase activity and telomere length. To test to contribution of these on C-circle production in LTT+ cells, we used a CRISPR/Cas9 approach to eliminate telomerase activity. We electroporated cells with dual *TERC* or a negative control (non-targeting) guide RNA-Cas9 ribonucleoproteins to establish four independent clonal lines from each treatment. We did this in cen3tel cells from a PD with long telomeres (PD171, 12 – 15 kb) and a PD with extremely long telomeres (PD643, 50 kb). The four independent *TERC* knockout (KO) clones each had a biallelic 84 bp deletion within the *TERC* pseudoknot domain and loss of hTR RNA (Figures S5A and S5B). As expected, the deletion completely ablated telomerase activity in each of the *TERC* KO clones (Figure 5A) and telomeres shortened progressively with propagation (Figure S5C). This later result confirmed that, despite the presence of C-circles in the parental lines, the cen3tel cells utilized telomerase rather than ALT as the TMM.

**Figure 5.**
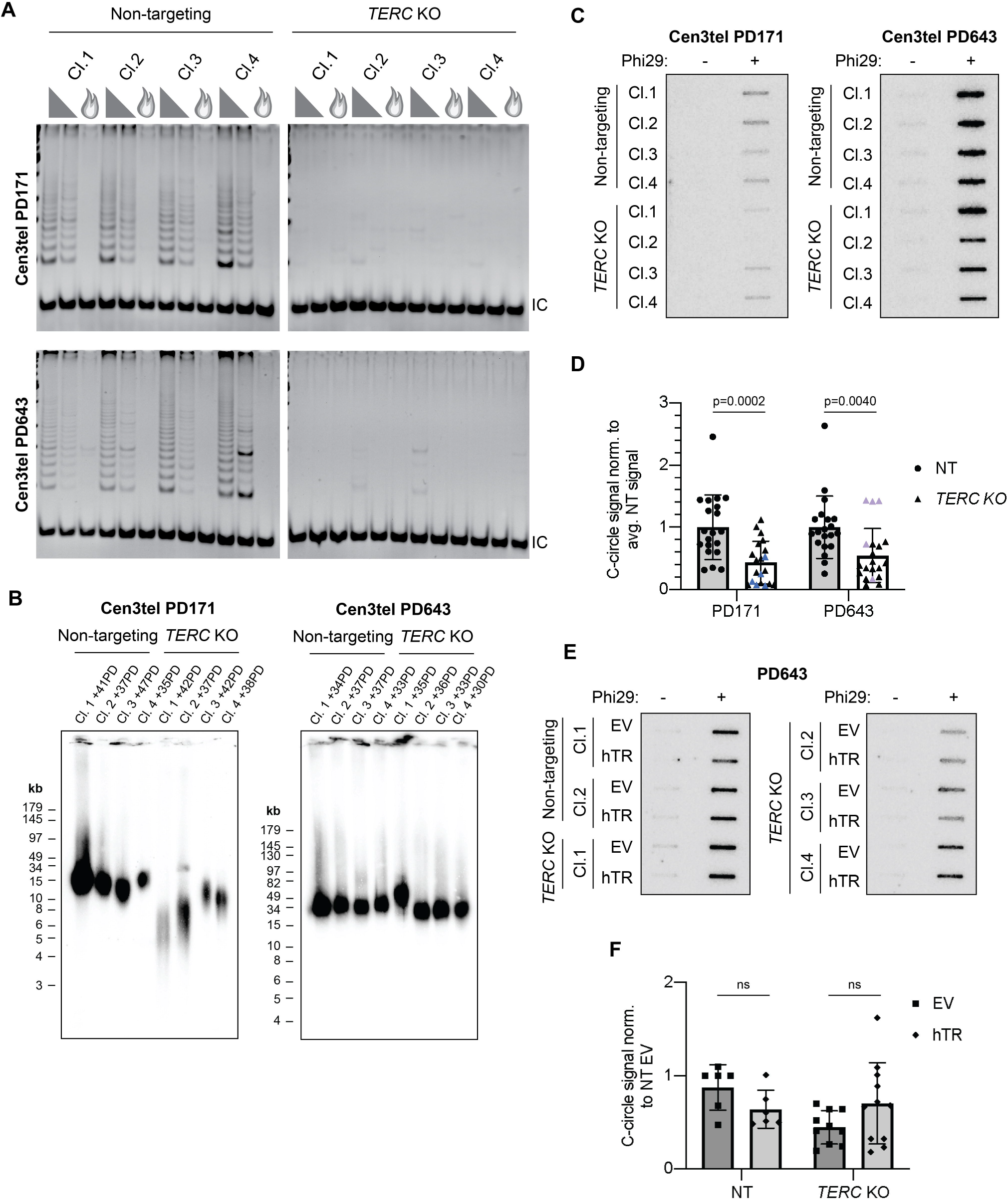
Telomerase activity does not contribute directly to C-circle production in cen3tel cells. (A) TRAP assay of cen3tel clonal populations from PD171 and PD643 that were non-targeted (NT) or *TERC* knockout (KO). Protein concentrations used in assays as in Figure 3A. IC, internal amplification control. (B) TRF analysis of the non-targeting and *TERC* KO subclones as in Figure 1. The analyses were performed at PDs post-clonal isolation that were similar to those of cells collected for CCAs. The differences in telomere length in the individual clones reflects the heterogeneity of telomere lengths in the parental polyclonal populations (Figure 1A). (C) CCAs of the same clonal populations. (D) Quantification of CCAs. The signal from each clone was normalized to the average signal of the non-targeting clones per replicate. For each, four samples were assayed over five experiments. Blue triangles, relative C-circle signals of PD171 *TERC* KO clone 1. Purple triangles, relative C-circle signals of PD643 *TERC* KO clone 1. Average +/− 1 standard deviation (SD). Statistical analysis was performed using an unpaired Student’s t-test. (E) CCA of cen3tel PD643 non-targeting or TERC KO clones that were transfected with empty vector (EV) or hTR-expressing plasmid. hTR expression and telomerase activity shown in Figure S6. (F) Quantification of CCAs as in (D), normalized to NT clones EV signal. Each sample assayed over three experiments, and outliers removed by interquartile test. Average +/− 1 SD. Statistical analysis was performed using an unpaired Student’s t-test. See also Figure S5–S7.

To assess the contribution of telomerase *per se* to C-circle production (e.g., by impacting repair at stalled or collapsed replication forks), CCAs were performed with cells that were collected as soon as possible after clone isolation to limit the extent of telomere attrition in the *TERC* KO clones and telomere elongation in the non-targeting clones. Although the amount of shortening or elongation could not be determined given each clone was derived independently from a polyclonal population with heterogenous telomere lengths, the lengths of most of the cen3tel *TERC* KO clones were shorter than the lengths of the non-targeting clones at PDs around when CCAs were performed (Figure 5B). The one exception was cen3tel PD643 *TERC* KO clone 1. However, its propagation for an additional 100 PDs demonstrated progressive shortening as expected with loss of telomerase-dependent TMM (Figure S5C).

We observed the average C-circle signal in the *TERC* KO clones was 50% of that in the non-targeting clones at both cen3tel PDs (Figures 5C and 5D). We noted, however, that the shortest PD171 *TERC* KO clone, clone 1 (blue triangle symbol), generally had the lowest, and the longest PD643 *TERC* KO clone, clone 1 (purple triangle symbol), generally had the highest C-circle signals relative to the other KO clones on repeated testing. Therefore, to investigate whether the lower average C-circle signal in the *TERC* KO clones was due to a secondary effect of telomerase deficiency on telomere length rather than the loss of a direct contribution of telomerase on C-circle production, we performed telomerase rescue experiments, transiently expressing hTR for 48 hr followed by CCAs. The 48 hr frame was much shorter than the 6 weeks required to isolate non-targeting and *TERC* KO clonal populations and propagate cells to a workable number for experiments; thus, any telomere lengthening as a result of restoration hTR expression and telomerase activity (Figures S6A and S6B) would be minimal. Moreover, we performed these experiments in *TERC* KO clones at cen3tel PD643, such that any telomere lengthening would be proportionally negligible to the starting telomere length. We found the C-circle signal did not increase in cen3tel PD643 *TERC* KO clones upon reintroduction of hTR and restoration of telomerase activity as compared to cells transfected with empty vector (Figures 5E and 5F). Thus, we conclude that the reduction in C-circle signal in the *TERC* KO clones was more likely due to telomere attrition than the loss of a direct effect of telomerase on C-circle formation.

We similarly ablated telomerase activity in the HeLaE1 cell line via *TERC* KO (Figures S7A– S7C). The HeLa E1 *TERC* KO clones had a narrower distribution of telomere lengths than three of the four non-targeting clones with loss of the longer telomeres (Figure S7D). In contrast to the cen3tel cells, loss of telomerase did not affect C-circle levels in HeLaE1 cells (Figures S7E and S7F). While HeLaE1 *TERC* KO clones appeared to have less high molecular weight telomere signal compared to non-targeting clones (Figure S7D), it is possible that the extent of telomere shortening and retention of very long telomeres were not sufficient to decrease C-circle signal in contrast to the cen3tel cells.

### LTT+ cells are variably sensitive to induced replication stress and FANCM does not restrain C-circle production

We next investigated whether the relationship between C-circle production and telomere length in the LTT+ cells was a result of increased replication stress. Telomeres are prone to replication fork stalling given the potential for secondary structures of G-rich repeats (36); thus, longer telomeres have greater opportunity for problematic replication. Replication stress is increased at ALT+ telomeres, making ALT+ cells exquisitely sensitive to agents such as HU, which induces replication stress through dNTP depletion. Therefore, we examined the sensitivity of LTT+ cells to HU-induced replication stress, exposing them to increasing concentrations of HU for 20 hrs. Then, nuclei were stained with Incucyte® Nuclight Rapid Red Dye, and cell proliferation was assessed by Incucyte® live cell imaging and nuclei counting over 96 hrs (Figure 6A). G292 cells were used as the ALT+ control for this experiment because U2OS nuclei did not stain with the dye. While G292 telomeres are shorter than U2OS telomeres (~20 kb compared to ~50 kb) (37), we found, as expected for an ALT+ cell line, G292 cells had C-circles (Figure 6B) and were very sensitive to HU, displaying impaired proliferation even at the lowest concentration tested (Figure 6A). Despite having cECTR signal in 2DGs and CCAs (Figure 2A and 2C), cen3tel PD185 cells, with average telomere length of 12 – 15 kb, were unaffected by HU exposure. Cen3tel PD650 cells, with telomeres of ~50 kb, exhibited diminished proliferation at the two higher concentrations of HU but not the low dose that impaired G292 cell proliferation. Lastly, both HeLa1.3 and HeLaE1 cells displayed reduced proliferation only at the highest HU concentration. Overall, the LTT+ cells exhibited a range of no sensitivity (cen3tel PD185) to some sensitivity to HU, though none was as severely affected as the G292 cells. This suggests that, while long telomeres may increase endogenous replication stress, ALT status may be a stronger factor.

**Figure 6.**
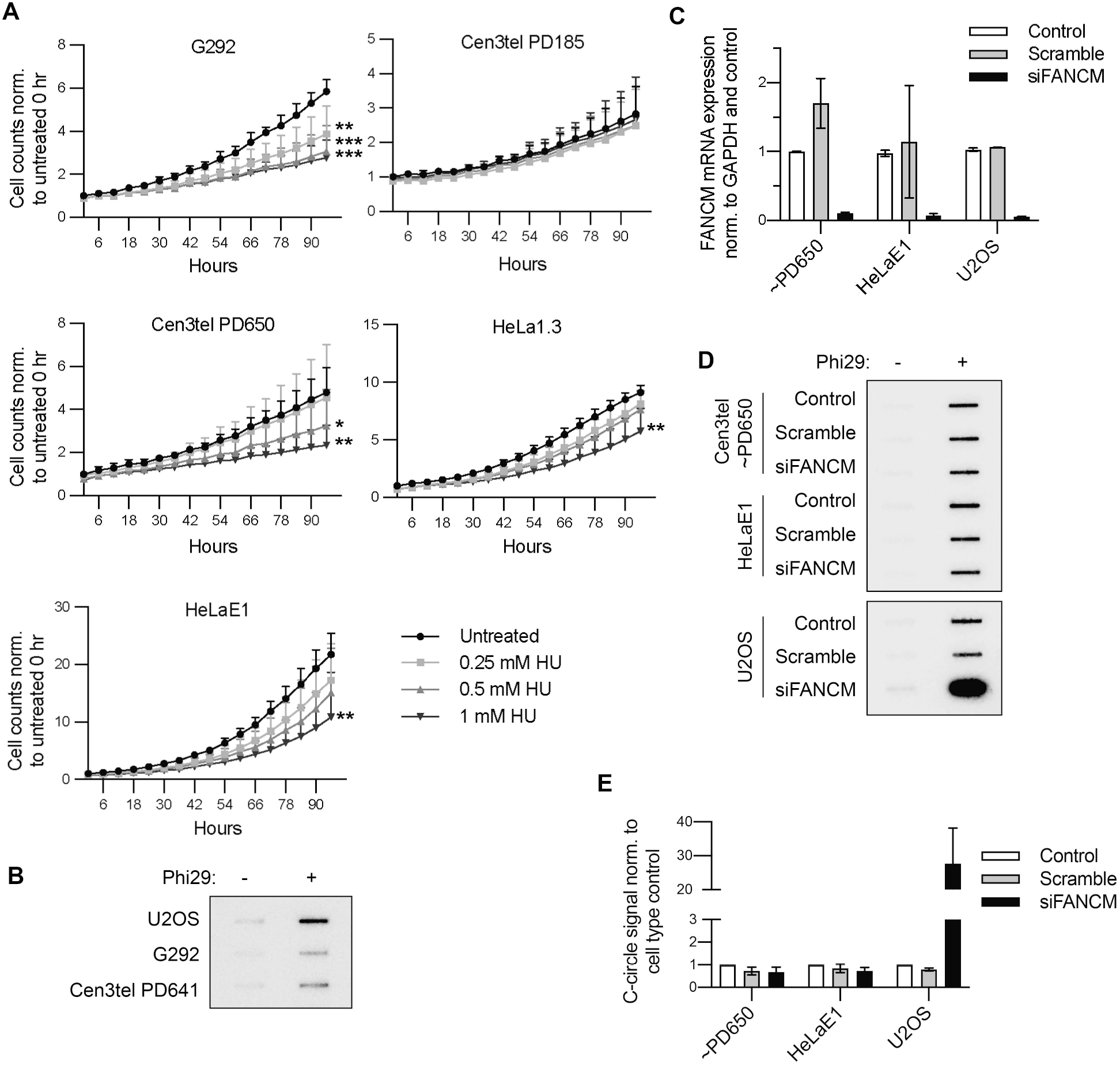
LTT+ cells experience some sensitivity to induced replication stress, though FANCM does not affect C-circle levels. (A) Average fold change in cell counts in the designated lines grown in the absence or presence of varying concentrations of hydroxyurea assessed by live cell imaging and automated nuclei counting over 96 hrs, then normalizing each count to the cell type’s average starting number of untreated cells. Each sample was performed in triplicate per experiment, and the experiment was performed twice. *=p≤0.05, **=0.001<p≤0.01, ***=p≤0.001. (B) Representative CCA displaying G292 C-circle levels compared to those of U2OS and cen3tel PD641 cells. N=3. (C) FANCM expression levels after FANCM or scramble siRNA knockdown assessed by RT-qPCR of FANCM normalized to GAPDH. Relative expression was normalized to the respective cell type’s untransfected control sample. N=2. (D) CCAs of cells from (C). (E) Quantification of CCAs as in (D). Amplification was performed in duplicate. Signal was normalized to the respective cell type’s average control signal. N=2.

FANCM assists with replication restart at collapsed replication forks and, thereby, functions as a repressor of C-circle production in ALT+ cells (26,27).Therefore, we assayed whether the FANCM protein affected C-circle production in LTT+ cells. We reduced FANCM expression through siRNA-mediated knockdown using two siRNAs to levels 5 – 10% of respective untreated control cells (Figure 6C). Whereas there was a marked increase in C-circle levels after FANCM knockdown in U2OS cells, there was no change in C-circle levels in either cen3tel PD650 or HeLaE1 cells (Figures 6D and 6E). These results reveal that FANCM is not involved in C-circle regulation in these LTT+ lines.

## DISCUSSION

Since their discovery, C-circles have been perceived to be specific to ALT+ cancers and mutually exclusive with telomerase activity (18). Prior reports of primary, stem, and immortalized cell lines have shown rare exceptions to this rule, though the C-circle levels were always markedly reduced compared to those of ALT+ lines (24,33). Here, we have presented three cell lines with robust telomerase activity that also displayed prominent C-circle signal, with the cells with the longest telomeres having levels comparable to that of ALT+ U2OS cells (Figures 2C and 3B). Additionally, analysis of cen3tel clonal populations still revealed telomerase activity and C-circle signal, consistent with co-occurrence in the same cell (Figure 3) and, thus, further evidence for a lack of mutual exclusivity.

While we did not directly assess ALT-dependent telomere synthesis in the LTT+ lines, several lines of evidence argue against this occurring. Previous work showed that APBs, which are the site for ALT-mediated synthesis (12), are absent or negligible in the cen3tel and HeLa1.3 cells, respectively (26,31,38). Additionally, the LTT+ lines had TRF profiles that lacked short telomere signal (Figure 1A and 4A) unlike ALT+ cells. Lastly, we found telomerase activity was necessary for telomere length homeostasis in the cen3tel cells (Figure S5C). Together, these findings underscore that C-circles are not exclusive to ALT+ cells but may also be a common feature of LTT+ cells. We also found that, in the context of hyper-elongated telomeres, C-circles were still produced in cells in which telomerase activity was ablated. Thus, CCAs as a screen for ALT cancers may lead to misidentification of tumors as ALT+ if they have very long telomeres, whether telomerase expressing or not.

### The mechanism of C-circle production is likely different between LTT+ and ALT+ cell lines

Given that C-circles are prominent in the various LTT+ lines, how are they produced? By analyzing different PDs of cen3tel cells, we observed a direct correlation of C-circle signal with telomere length. Importantly, we also observed decreased C-circle signal after telomere attrition in cen3tel *TERC* KO clones, but this decrease was not rescued by short term telomerase re-expression (Figures 2, 5, S5, and S7). Thus, our assessments provide thorough evidence that it is telomere length and not a direct involvement of telomerase that positively contributes to C-circle production in the LTT+ cells.

It has been proposed previously that increased telomere length in telomerase+ lines leads to C-circle production through heightened replication stress (24), which is also a mediator of C-circle production in ALT+ cells (30,39). However, distinct from ALT+ cells (23,26–28,39,40), we found FANCM expression did not negatively regulate C-circles in the LTT+ cells (Figures 6C–6E). This strongly suggests that replication stress is not a required contributor to C-circle production. Consistent with this, cen3tel PD185 demonstrated no sensitivity to HU (Figure 6A) yet had detectable cECTRs in both 2DGs and CCAs (Figures 2A and 2C). Thus, the mechanism(s) regulating C-circle production in these LTT+ lines appears distinct, at least in one respect, from that of ALT+ cells. Future investigations could assess whether other factors associated with C-circle production in ALT+ cells contribute to C-circle production in LTT+ lines or if alternative factors are involved.

### cECTR assays are not interchangeable

Our data underscore differences in the detection of telomeric circles by the three main cECTR assays. The TCA is frequently used as a substitute for 2DG electrophoresis despite having the capacity to detect fewer types of cECTRs due to the requirement of an intact strand (17). Although typically used interchangeably, the 2DGs did, and the TCAs did not, yield signal for the LTT+ cells (Figures 2 and 4). Moreover, we observed a prominent signal in CCAs while cECTR amplification in the TCA was absent. The detection of a cECTR arc in 2DGs but no signal in TCAs can be explained by nicks or gaps on both strands of double-stranded telomeric circles or the arc solely representing looped telomeric fragments. However, the detection of cECTRs in CCAs but not TCAs is unexpected because, theoretically, the C-circle molecule should be amplified in both assays (when the TCA specifies the C-rich template). The results obtained in the LTT+ lines could reflect a combination of biological factors and technical aspects. The biological factor could be that C-circles are at a low abundance and other cECTRs are absent, and the technical aspect could be that the CCA is more sensitive than the TCA as previously demonstrated (41), enabling it to amplify the low abundance C-circles. ALT+ cells may contain various cECTR molecules so that, even if at low abundance, together they allow for detection by both assays. Future investigations assessing the structure and abundance of cECTRs in LTT+ and ALT+ cell lines will be helpful for understanding differences of detection.

### Continuous telomere elongation in the absence of t-circle production

The absence of t-circles as detected in TCAs in the LTT+ lines supports the association of these cECTRs specifically with telomere trimming to achieve telomere length homeostasis. Similar to telomeres in the cen3tel cells, telomeres in the HeLaE1 cells continuously lengthened across propagations (31,35). The telomere lengthening in HeLaE1 cells was dependent on overexpression of both TERT and hTR, which was proposed to result in the telomeres being maintained in an extendable state (31,35). Though its copy number was not experimentally changed, hTR levels in the cen3tel cells increased 3.5-fold at PD30 compared to fibroblast hTR levels, which then returned to normal levels before increasing gradually to 4.5-fold when telomeres were 100 kb (31). Thus, telomerase overexpression in the LTT+ lines may not only be sufficiently high to maintain the telomere in an extendable state but also prevent telomere trimming, thereby overriding its contribution to telomere length homeostasis.

## EXPERIMENTAL PROCEDURES

### Cell Lines, Culturing, and Handling

Cell lines used in this study include: cen3tel (31), HeLa1.3, HeLaE1, U2OS, G292, 293T, BJ fibroblasts, and BJtert. Cen3tel (female) and HeLa1.3 (female) cells were cultured in DMEM supplemented with 10% FBS, 1X non-essential amino acids, and 1X GlutaMAX (ThermoFisher). HeLaE1 (female) cells were grown in DMEM +10% FBS. U2OS (female) and G292 (female) cells were cultured in McCoy’s 5A +10% FBS. 293T cells were grown in MEM-alpha +10% FBS. BJ fibroblast (male) cells were grown in DMEM +10% FBS, and BJtert (male) cells were grown in a 4:1 mix of DMEM and Medium 199 +10% FBS. Population doublings were added based on how much the cells were split at each passage (i.e., cells are split 1:8, +3PDs). All cells were grown at 37°C and 5% CO_2_. Cell lines were authenticated every 6 – 12 months through STR profiling at MD Anderson’s Cytogenetics and Cell Authentication Core and were routinely checked for mycoplasma (SouthernBiotech #13100-01). Cen3tel cells were authenticated as having distinct STR profiles from all other cell lines and being similar to each other across PDs. When clonal populations were necessary, IDT’s array dilution method was employed to isolate them. Briefly, 4,000 cells were plated in a single well of a 96-well plate. These were diluted 1:1 vertically down the column, then diluted 1:1 horizontally across all rows. When a single clone was visibly growing in a well, it was picked and transferred to a larger plate.

### *TERC* CRISPR/Cas9 Knockout

*TERC* was knocked out via ribonucleoprotein (RNP) electroporation of Cas9 protein with two guide RNAs. The guides were designed by Horizon’s CRISPR Design Tool, where crRNA is the target and tracrRNA (cat. #: U-0020005-05) is the scaffold. The catalogue numbers for the *TERC* guides are: crRNA-539475:WCEFR-000001 and crRNA-539476:WCEFR-000002. A non-targeting crRNA was also used as a negative control in each experiment (cat #: U-007501-01-05). To prepare the RNP, 1μL of 200 μM crRNA was mixed with 1 μL of 200 μM tracrRNA. Then 1.6 μL 10 mg/mL Cas9 protein (IDT cat. #: 1081059) was added to that mixture and incubated at room temperature for 10-15 min. This mixture was resuspended with 1×10^6^ cells in 100 μL of Lonza’s Nucleofector Solution Kit V and electroporated in an Amaxa electroporator on setting D-032. Cell mixtures were then transferred to a 6-well plate and allowed to recover 24 – 48 hrs before assessing their genotypes.

### siRNA Transfection

Sequences for FANCM DsiRNA were obtained from (27) (labeled siFa and siFb in that paper). DsiRNA’s were custom ordered from IDT and resuspended in nuclease-free water. Cells were seeded in 6-well plates so that the confluency 24 hr later would be ~60%. Both DsiRNAs against FANCM were mixed with 5 μL Lipofectamine RNAiMAX (ThermoFisher cat #: 13778150) and 500 μL Opti-MEM so that the total final concentration of RNA in media would be 40 nM. The mixture was incubated at room temperature for 5 min. Scramble siRNA was utilized as a negative control in each experiment at the same concentration (Sigma-Aldrich cat. #: SIC001). This siRNA mixture was added to 1.5 mL fresh complete medium on the cells. Cells were collected 48 hrs after knockdown, half to be analyzed for FANCM mRNA expression by RT-qPCR, and the other half for C-Circle Assay.

### hTR Overexpression

Cen3tel PD643 NT and *TERC* KO clones were transfected with 2 μg empty vector (pcDNA3.1+) or hTR-expressing plasmid (pBS-U1-hTR). A pellet of 1×10^6^ cells was resuspended in 100 μL Lonza’s Nucleofector Solution Kit V and electroporated in an Amaxa electroporator on setting D-032. Transfected cells were then transferred to a 6-well plate, media was replaced with fresh media 24 hr later, and cells were collected an additional 24 hr later (48 hrs after electroporation) to assess hTR expression, telomerase activity, and/or C-circle signal.

### RT-qPCR

RNA was extracted from cell pellets using QIAGEN’s RNeasy Mini Kit (cat. #: 74106), following the kit’s protocol and performing the optional DNAse treatment. RNA concentration was determined by Nanodrop. Synthesis of cDNA was performed using Quantabio’s qScript cDNA Synthesis Kit (cat. #: 95047), amplified with random hexamers for hTR and oligo-dT primers for FANCM assays and the kit’s prescribed conditions. RNA (500 – 1000 ng) was reverse transcribed in a 20 μL reaction with 1 μL qScript RTase for 90 min at 42°C. The cDNA was then diluted to 5 ng/μL in RNAse-free H_2_O. For qPCR, Applied Biosystems’ PowerUp™ SYBR™ Green Master Mix was used (cat. #: A25742), and amplification occurred with 250 nM each of forward and reverse primers (sequences below). Applied Biosystems’ QuantStudio™ 6 Flex Real-Time PCR System was used to determine CT values of amplified samples (performed in technical triplicates). The volume per reaction was 20 μL, and 4 μL of that was cDNA. The qPCR amplification conditions were were 50°C for 2 min, 95°C for 2 min, and 40 cycles of 95°C for 15 sec and 60°C for 1 min. The hTR amplicon was 126 bp, the FANCM amplicon was 118 bp, and the GAPDH amplicon was 193 bp. Specificity of the primers was verified by an *in silico* screen (NCBI BLAST). CT values that were greater than 0.05 away from the next closest value in a set of triplicates were considered outliers and not included in the analysis. Values were normalized for loading by subtracting the respective GAPDH CT values from the FANCM CT values for each sample, creating ΔCT values. Then they were normalized to the respective untreated sample for each cell line, creating ΔΔCT values. The fold change of each sample was determined (RQ value) after inversing the logarithm, and the averages of these numbers were plotted.

hTR forward primer: TCTAACCCTAACTGAGAAGGGCGTAG

hTR reverse primer: GTTTGCTCTAGAATGAACGGTGGAAG

FANCM forward primer: AGCGCAGATTTCCTATAAACAGA

FANCM reverse primer: CCTCTTCTGGCATTCCCGTT

GAPDH forward primer: CACATGGCCTCCAAGGAGTAAG

GAPDH reverse primer: TACATGACAAGGTGCGGCTCCC

### Telomeric Repeat Amplification Protocol

TRAP assays were performed using Sigma-Aldrich’s TRAPeze Telomerase Detection Kit (cat. #: S7700) non-radioactive detection by staining method with some modifications. CHAPS buffer was supplemented with 1:100 protease inhibitor (Millipore cat. #: 539134) and 1:400 RNase inhibitor (Promega cat. #: N251B). Protein lysates were serially diluted to concentrations of 0.6 μg/μL, 0.03 μg/μL, and/or 0.006 μg/μL, and 1 μL from these was added to each reaction. Final protein concentrations per experiment are listed in the figures and figure legends. Amplification reactions were 25 μL total (using half the volumes described by the kit). Each reaction received 1 unit Titanium Taq polymerase (TaKaRa cat. #: 638517). All steps prior to amplification were performed RNase-free. Amplification conditions were: 30°C for 30 min; 95°C for 2 min; 34 cycles of 94°C for 15 sec, 59°C for 30 sec, 72°C for 1 min; and 4°C. Total products were loaded in Bio-Rad vertical plate gels (12.5% 19:1 acrylamide/bis, 0.5X TBE, 0.1% ammonium persulfate, TEMED) for 100 V over 2.5 hr in 0.5X TBE. Gels were stained in 1:10,000 ethidium bromide diluted in ddH_2_O for 30 min, washed in ddH_2_O, and imaged using a ChemiDoc-ItTS2 Imager, UVP.

### C-Circle Assay

Amplification of C-circles was performed as described in (42) with minor modifications. QCP lysates were constantly stored on ice when setting up reactions. DNA concentrations were determined using Invitrogen’s Qubit 4 Fluorometer. DTT was added fresh to each reaction. Samples were diluted to 10 – 40 ng/μL DNA in QCP buffer, and 1 μL was added per reaction. Amplification conditions were: 8 hr at 30°C, 20 min at 70°C, 10 min at 22°C. Each reaction included a negative control lacking phi29 polymerase to reveal background signal. Many assays were performed in technical duplicates. Products were loaded onto positively charged Zeta-probe membranes via slot blot. Membranes were pre-hybridized in Church buffer (1mM EDTA, 1% BSA, 0.5M phosphate buffer pH 7.2, 7% SDS) at 37°C for 30 min, then hybridized to 5×10^5^ cpm/mL ^32^P-end-labeled (CTAACC)_3_ probe overnight. Washes of blot after hybridization were performed flat and shaking at room temperature with wash buffer (0.5X SSC, 0.1% SDS), pre-warmed to 37°C, 4X for a total of 20 min. C-circle signal was exposed to a PhosphorImager screen overnight and detected on an Amersham Typhoon scanner. Quantification was performed by assessing signal intensity using Image Quant TL software, and technical duplicates were averaged. For *TERC* KO blots, the four non-targeting clones’ signals were averaged, then each clones’ signal (non-targeting and *TERC*) were normalized to that number for final analysis. Statistical significance was assessed by performing one-way ANOVA analyses of three biological replicates.

To determine whether the templates in the CCAs were circular, section 7.5 from (42) was followed with one modification to protect C-circles from shearing after exonuclease digest: 2 μg pcDNA3.1 plasmid were added before exonuclease treatment to mock and exonuclease samples (digested with 12.5 U lambda exonuclease and 100 U exonuclease I). CCA amplification was performed as described above. Exonuclease digest was assessed by agarose gel.

### Telomere Length Analysis

Pellets of 1×10^6^ cells were resuspended in a 1:1 mixture of 2% agarose and 1X PBS, then solidified in a plug mold. Plugs were digested in proteinase K buffer (100mM EDTA pH 8.0, 0.2% Na deoxycholate, 1% SDS, and 1mg/mL proteinase K) overnight at 50°C. Then, the plugs were washed at room temperature while rotating in 1X TE for 1 hr 3 times, 1X TE + 1 mM PMSF for 1 hr, 1X TE for 1 hr, ddH_2_O for 20 min, and 1X Cutsmart buffer for 1 hr. The plugs were digested overnight in 60 units HinfI and 60 units RsaI at 37°C. After washing in 1X TE for 1 hr and equilibrating in 0.5X TBE for 1 hr at room temperature, the plugs were loaded onto a 1% pulsed field (PFGE) certified agarose (Bio-Rad cat. #: 1620137) agarose gel. Electrophoresis was performed in a Bio-Rad CHEF-DR II system. A low molecular weight ladder (NEB cat. #: N3232L) and high molecular weight ladder (NEB cat. #: N0342S) were included in each length assessment. Running conditions for gels assessing cells with average telomeres (BJtert, cen3tel PDs < 600) were 14°C, 12 hrs, 6 V/cm, 5-20 sec switch time. Running conditions for gels assessing cells with long telomeres (HeLa1.3, HeLaE1, cen3tel PDs > 600) were 14°C, 16 hrs, 6 V/cm, 5 −20 sec switch time. After electrophoresis, gels were stained in 1:10,000 ethidium bromide and washed in depurination solution for 20 min (0.25M HCl), denaturation solution for 30 min (0.5M NaOH, 1.5M NaCl), and neutralization solution for 30 min (0.5M Tris pH 1.5, 3M NaCl), with brief ddH_2_O washes in between. DNA was transferred to positively-charged Zeta probe membrane. Blots were prehybridized in Church buffer (1mM EDTA, 1% BSA, 0.5M phosphate buffer pH 7.2, 7% SDS) at 65°C, rotating for 2 hr, then hybridized overnight to a ^32^P-labeled 800 nt C-rich telomeric fragment amplified by random labeling. The 800 bp telomere repeat fragment was digested from pSP73.Sty11 (telomere repeat-containing plasmid) with EcoRI and purified by gel extraction; labeling the C-rich strand was specified by synthesizing in the presence of dCTP isotope. Telomere signal was exposed to a PhosphorImager screen 4 hrs – overnight and detected on an Amersham Typhoon scanner.

### 2D Gel Electrophoresis

DNA was isolated using QIAGEN’s DNeasy Blood and Tissue Kit and digested with HinfI, RsaI, and RNaseA overnight at 37°C. Samples were phenol/chloroform purified and EtOH precipitated. For each sample, 5 – 10 μg DNA were loaded into a 0.4% agarose gel (10×12 cm) and electrophoresed for 15 hrs at 1 V/cm. After staining with ethidium bromide, entire lanes were excised from the gel and rotated 90° counterclockwise onto a 20×25 cm tray. The second dimensional gel was run in 1% agarose at 4°C for 8 hr at 5 V/cm. The remainder of the assay (gel washes, transfer, hybridization, and exposure) was performed similarly to the telomere length analysis.

### T-Circle Assay

Assays in Figures 2 and 4: Genomic DNA was extracted from pellets containing 1 million cells by resuspending in 150 μL of QCP lysis buffer (50 mM KCl, 10 mM Tris pH 8.5, 2 mM MgCl_2_, 0.5% NP40, 0.5% Tween20, 0.05 U/mL QIAGEN protease) and following the procedure described in (42). The annealing mix was prepared by adding 400 ng of genomic DNA to 10 mM Tris pH 8.0 in a final volume of 26 μL, followed by the addition of 10 μL of 10 μM TelC (CCCTAA)_4_ oligo and 4 μL of annealing buffer (200 mM Tris-HCl pH 8.0, 200 mM KCl, 1 mM EDTA). The 40 μL annealing mix was incubated in a thermocycler at 96°C for 5 min and brought gradually to 25°C using the ramp option (0.1°C down/sec). The annealing mix was divided into two 20 μL aliquots, and each of these was mixed with 18.5 μL of master mix (2X phi29 polymerase buffer, 0.4μM dNTP mix, 0.4 mg/ml BSA) and with 1.5 μL of either phi29 polymerase (15 U) or nuclease-free ddH_2_O. Primer extension was achieved by incubating the reaction at 30°C for 12 hr, and phi29 polymerase was then inactivated by incubation at 65°C for 20 min. The extension products were separated by gel electrophoresis (0.6% agarose in 1X TAE), by first running at 3.5 V/sec for 1 hr then decreasing the voltage to 1.3 V/sec for 18 hr. The gel was then subjected to depurination (10 min in 0.25 M HCl), denaturation (15 min in 1.5 M NaCl, 0.5 M NaOH, twice), and neutralization (15 min in 1.5 M NaCl, 0.5 M Tris pH 7.2, twice), and DNA was transferred onto a positively-charged nylon membrane. The DNA was crosslinked onto the membrane at a setting of 120 mJ/cm^2^, and the membrane was incubated with pre-hybridization buffer (5X SSC, 0.1% sarkosyl, 0.04% SDS) for 1-2 hours at 65°C. The t-circle extension products were finally detected by incubating the membrane with 1.3 nM of Digoxigenin-labeled TTAGGG-rich (DIG-TelG) probe (generated as explained in (43)) followed by detection of the DNA bound DIG-TelG probe as explained in (44).

Assays in Figure S2: DNA was extracted using QIAGEN’s DNeasy Blood & Tissue Kit, then digested overnight in either HinfI + RsaI or ExoV at 37°C. Samples were phenol/chloroform extracted and EtOH precipitated. For each, 500 ng DNA in 10 μL were mixed with 10 μL 2X annealing buffer (400 mM Tris pH 7.5, 400 mM KCl, 2 mM EDTA) and 50 pmol oligo ((CCCTAA)_4_ or (TTAGGG)_4_), then denatured at 95°C for 5 min. Mixtures were allowed to cool to room temperature for 30-60 min while annealing. Next, annealed samples were split in half, each receiving 10 μL amplification buffer (0.2 mM dNTPs, 0.2 mg/mL BSA, 1X phi29 buffer) with either 7.5 U phi29 polymerase or ddH_2_O. Samples were amplified in a thermocycler at 37°C for 12 hrs, and then the reaction was terminated by heating to 65°C for 20 min. Products were separated by running in a 0.5% agarose gel with ethidium bromide in 0.5X TBE first at 120V for 1 hr, then at 40 V for 16 hr. The gel washes, transfer to membrane, hybridization, and exposure were performed similarly to what is described in telomere length analysis. Hybridization was performed with either a ^32^P-labeled 800 nt C-rich or G-rich telomeric fragment amplified by random labeling, corresponding to which oligo was used for annealing.

### Assessment of Proliferation

Cell lines were plated in 96 well plates. The number of cells plated varied based on the cell type and was determined by respective proliferations in a pilot assay. Hydroxyurea (diluted in ddH_2_O) was added to final concentrations of 0.25, 0.5, and 1 mM, and some cells were left untreated as a negative control. Samples were performed in triplicate. After incubating for 20 hr in HU, the media was replaced with fresh media, including Incucyte® Nuclight Rapid Red Dye (Sartorius, cat. #: 4717) diluted 1:2000. The plate was incubated at room temperature for 30 min before placing in the Incucyte® Instrument. Whole wells were imaged every 6 hrs for a total of 96 hrs. Counts of nuclei per well at each time point were assessed by Incucyte® Analysis Software (v2020C). Further analysis was performed manually: the average number of cells at time 0 for the untreated samples was determined, then each subsequent count was normalized to that number. Normalized values were plotted to observe the effect on proliferation over time. Statistical analyses were performed by comparing the HU samples to the untreated sample within a cell type at the final time point (96 hrs) using a Student’s t-test.

### Statistical Analyses

Statistical analyses were performed on GraphPad Prism software (version 8.0). Two-tailed unpaired Student’s t-test and two-way analysis of variance (ANOVA) were used for statistical analyses, as noted in the corresponding figure legend. The number of biological replicates for each experiment is noted in the corresponding figure legend, and averages are listed as the mean +/− standard deviation. Outliers were omitted according to the interquartile range test. Statistical significance was defined as p<0.05. GraphPad Prism was used to create the graphs.

## DATA AVAILABILITY

All data are contained within this manuscript. Cell lines generated in this study are available upon request from Dr. Alison Bertuch.

## SUPPORTING INFORMATION

This article contains supporting information.

## ACKNOWLEDGEMENTS

We thank Dr. Titia de Lange (The Rockefeller University) and Dr. Joachim Lingner (École Polytechnique Fédérale de Lausanne) for generously sending us the HeLa1.3 and HeLaE1 cell lines, respectively. We also thank Dr. Fabio Stossi and Ms. Hannah Johnson at Baylor College of Medicine’s Integrated Microscopy Core for their assistance in using the Incucyte® Instrument.

## AUTHOR CONTRIBUTIONS

Conceptualization, C.Y.J. and A.A.B; Investigation, C.Y.J., C.L.W., S.P.M., and D.K.M.; Resources, C.M.; Writing – original draft, C.Y.J., Writing – reviewing and editing, C.L.W., S.P.M., D.K.M., C.M., J.K., and A.A.B.; Supervision, J.K. and A.A.B.; Project administration, A.A.B.; Funding acquisition, A.A.B.

## FUNDING AND ADDITIONAL INFORMATION

This work was supported by the National Institutes of Health [T32GM08307 to C.Y.J., R01CA211653 and R01GM07750 to A.A.B, and R01CA227934, R01CA234047, R01CA228211, and R01AG077324 to J.K.]; and the Human Frontier Science Program [LT000108/2019-L to S.P.M.]. The content is solely the responsibility of the authors and does not necessarily represent the official views of the National Institutes of Health.

## CONFLICT OF INTEREST

The authors declare that they have no conflicts of interest with the contents of this article.

## ABBREVIATIONS

2DG: 2 dimensional gel
ABPs: ALT-associated PML bodies
ALT: alternative lengthening of telomeres
BJtert: TERT-immortalized BJ
CCA: C-circle assay
cECTRs: circular extrachromosomal repeats
hESCs: human embryonic stem cells
HU: hydroxyurea
KO: knockout
LTT+: long telomere, telomerase +
PDs: population doublings
PML: promyelocytic leukemia
t-circles: telomeric circles
TCA: t-circle assay
TMM: telomere maintenance mechanism

## Notes

### Competing Interest Statement

The authors have declared no competing interest.

