## Supporting Information for "Hyperextended telomeres promote C-circle formation in telomerase positive human cells"

Supplementary Figures 1 – 7

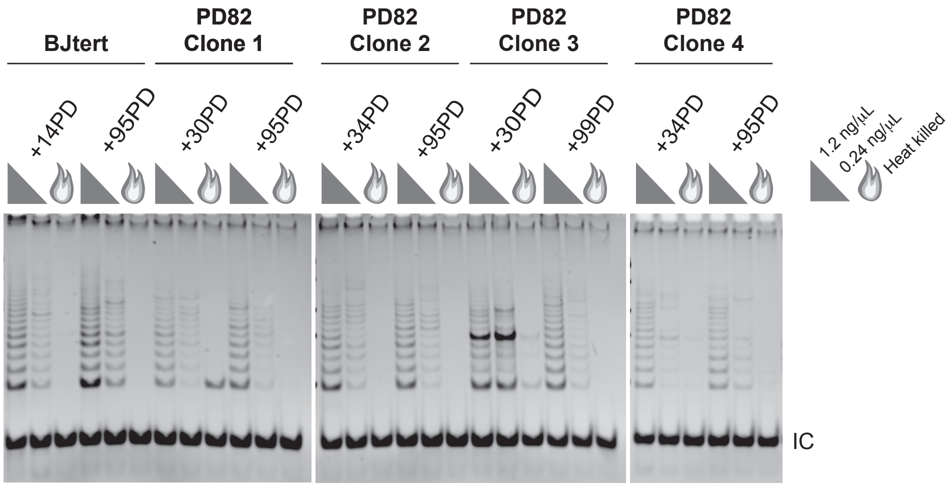


**Figure S1. Telomerase activity in BJtert cells is similar to that of cen3tel PD82 clones** TRAP assays of BJtert cells and clonal populations of cen3tel PD82 cells at subsequent early and late PDs. IC, internal amplification control.

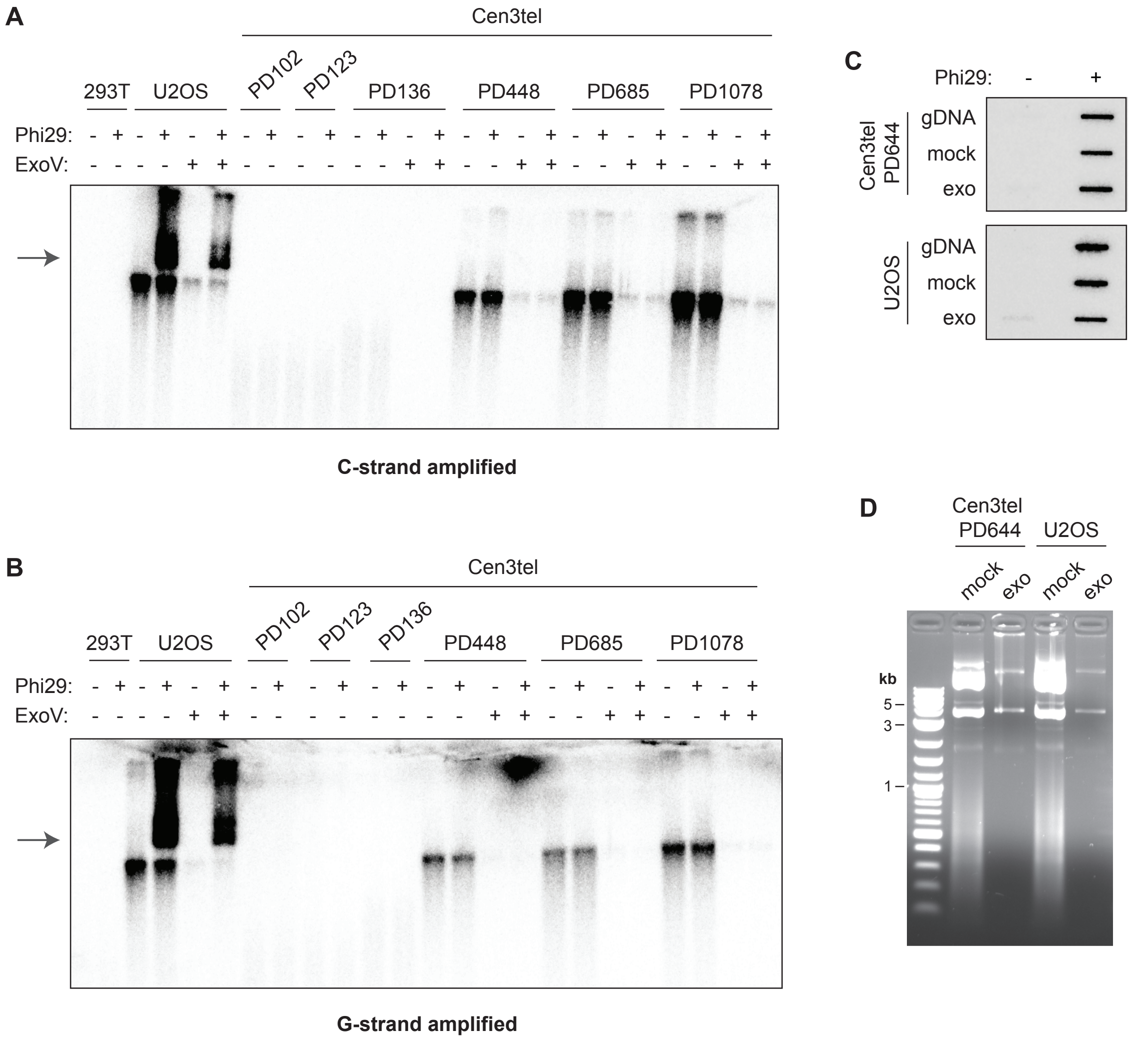


**Figure S2. cECTRs are not detected in cen3tel cells by t-circle assay (TCA), irrespective of which strand is amplified** (A) cECTR analysis by TCAs of the designated cells using a G-rich primer and a radiolabeled C-rich telomere probe to detect the amplified product. Arrow denotes amplified cECTR product. ExoV was used to degrade non-circular DNA prior to TCAs. (B) TCA as in (A) except using a C-rich primer and radiolabeled G-rich telomere probe to detect the amplification product. The same DNA was used as in part (A). (C) Cen3tel PD644 and U2OS DNA were extracted by Qiagen DNeasy kit and restriction digested to fragment genomic DNA. Then, after spiking with pcDNA3.1 (which lacks telomeric repeats), DNA samples were digested with lambda exonuclease and E. coli exonuclease I (exo sample) or no enzyme (mock sample). CCAs were then performed as normal. gDNA is genomic DNA that was never digested. (D) 1% agarose gel of the exonuclease treated samples. Samples same as in (C). The first lane is NEB 1 kb ladder.

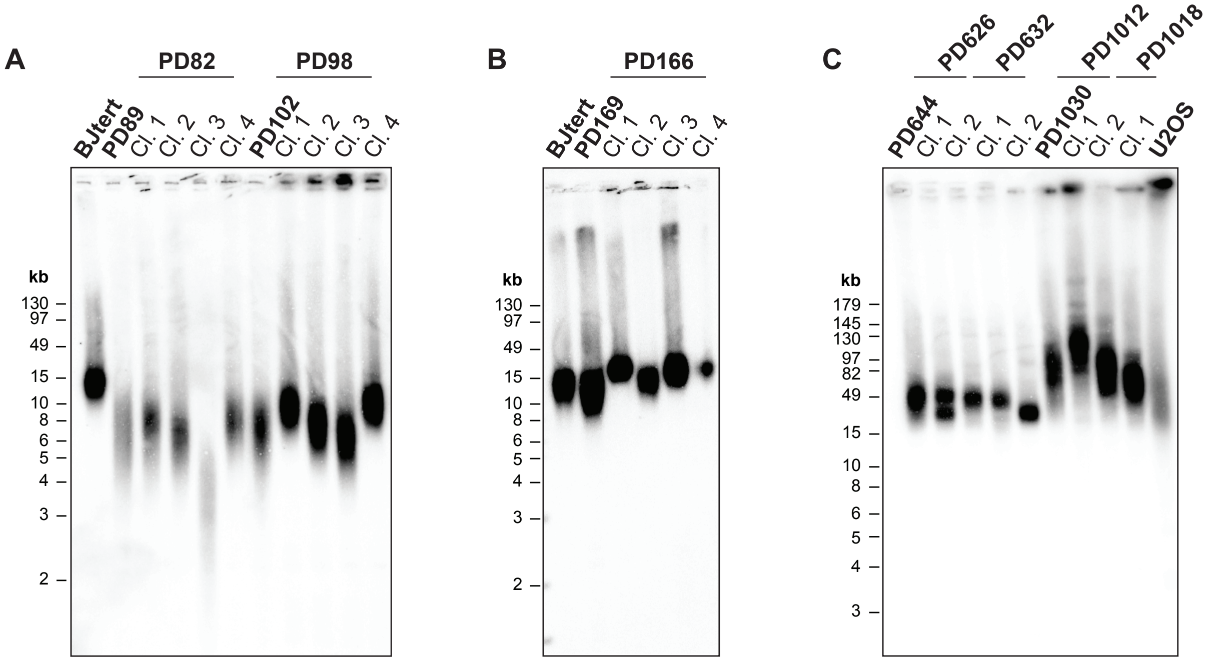


**Figure S3. Telomere lengths in cen3tel clonal populations** (A, B, C) TRF analysis was performed as in Figure 1. Telomere lengths of clones were compared to those of polyclonal populations of similar PDs.

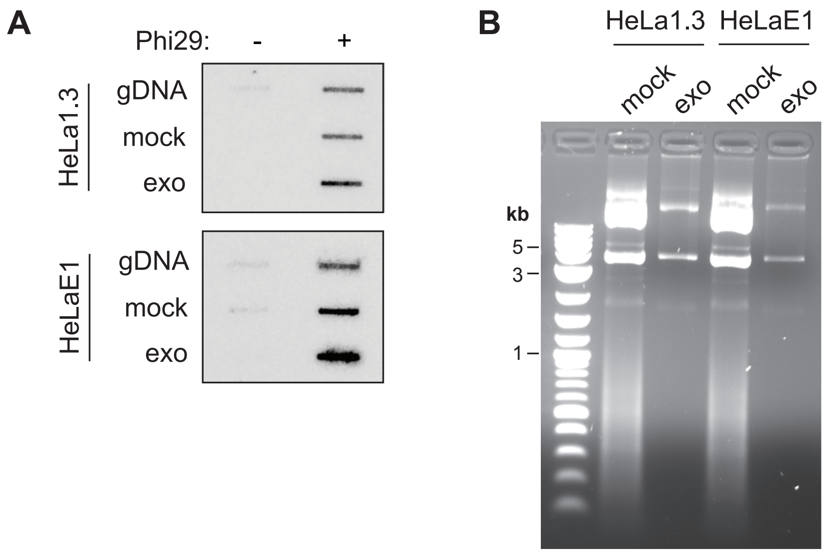


**Figure S4. The template amplified in the HeLa1.3 and HeLaE1 cells CCA assays is circular** (A) Same as in Figure S2C. DNA from HeLa1.3 and HeLaE1 cells. (B) Same as in Figure S2D, with the same DNA from (A).

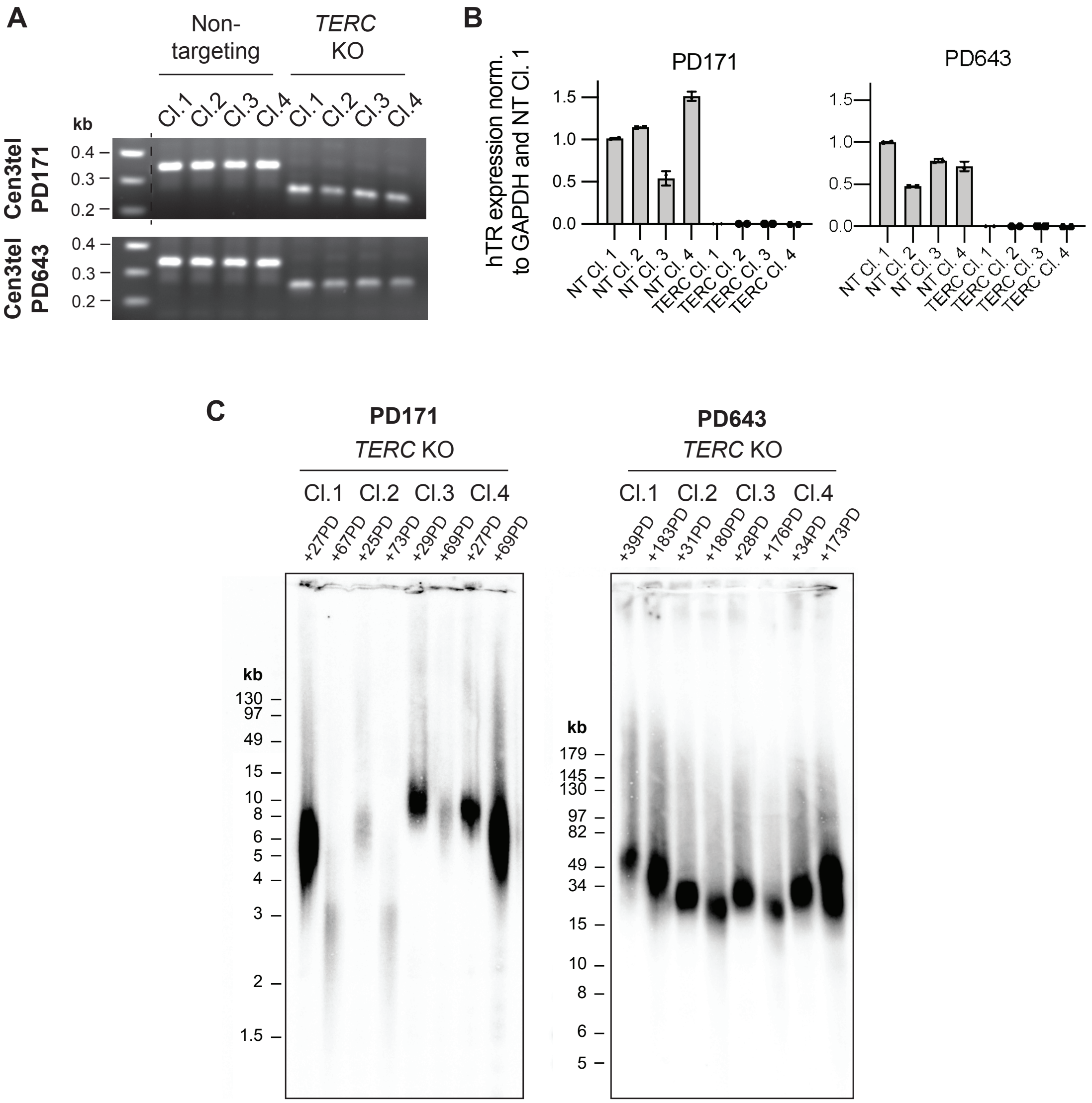


**Figure S5. *TERC* gene deletion ablates hTR expression and telomerase activity and leads to telomere shortening in cen3tel PD171 and PD673 cells** (A) PCR amplification of the *TERC* genomic locus to assess the 84 bp deletion. The 337 bp band reflects WT sequence, and the 253 bp band reflects *TERC* KO. (B) RT-qPCR assessment of hTR expression in non-targeting (NT) and *TERC* KO clones. Levels were normalized to GAPDH expression and NT clone 1. (C) TRF analysis of non-targeting and *TERC* KO subclones as in Figure 1 to assess telomere attrition between an early and late collection for each clone.

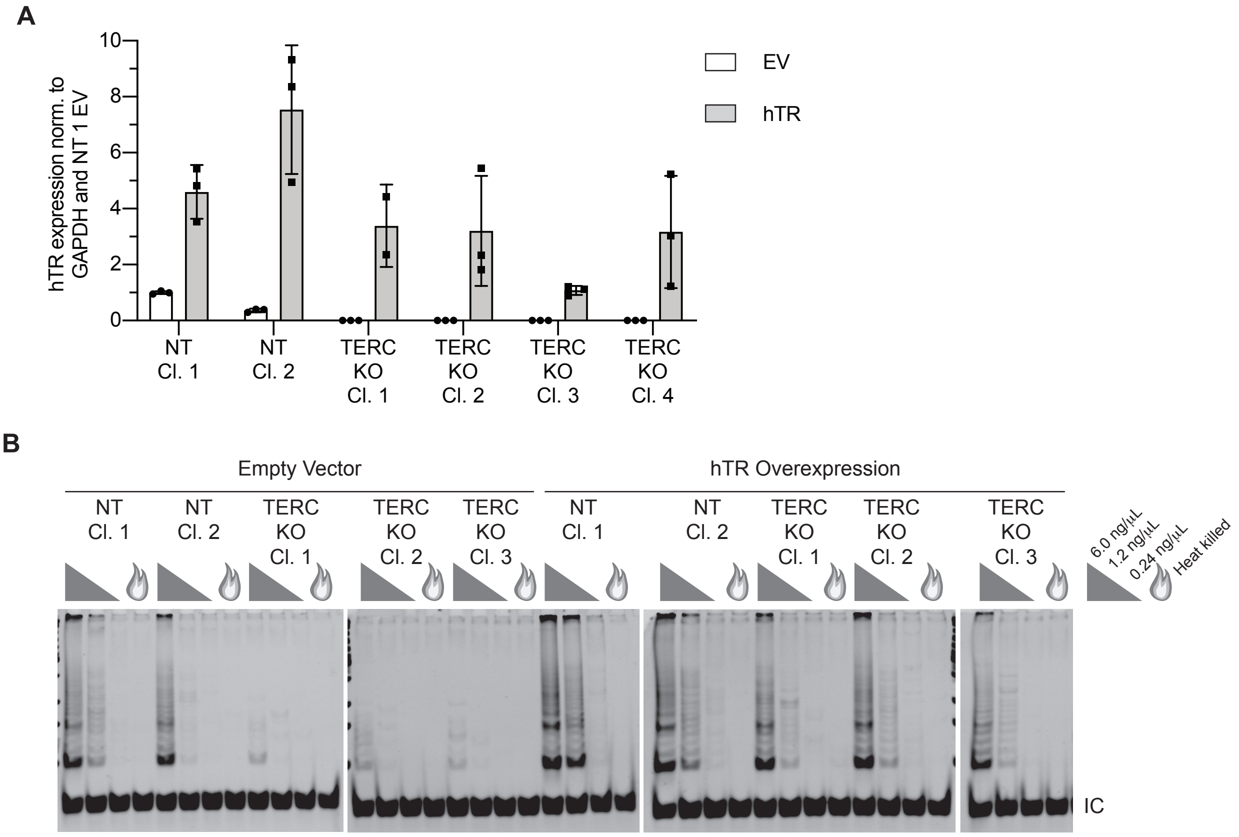


**Figure S6. hTR overexpression restores telomerase activity in *TERC* KO clones** (A) RT-qPCR assessment of hTR expression after transfection with empty vector (EV) or hTR-containing plasmid in non-targeting (NT) and *TERC* KO clones. Levels were normalized to GAPDH expression and NT clone 1 EV. N=3. (B) TRAP assay of clones following the same treatment as in (A). IC, internal amplification control.

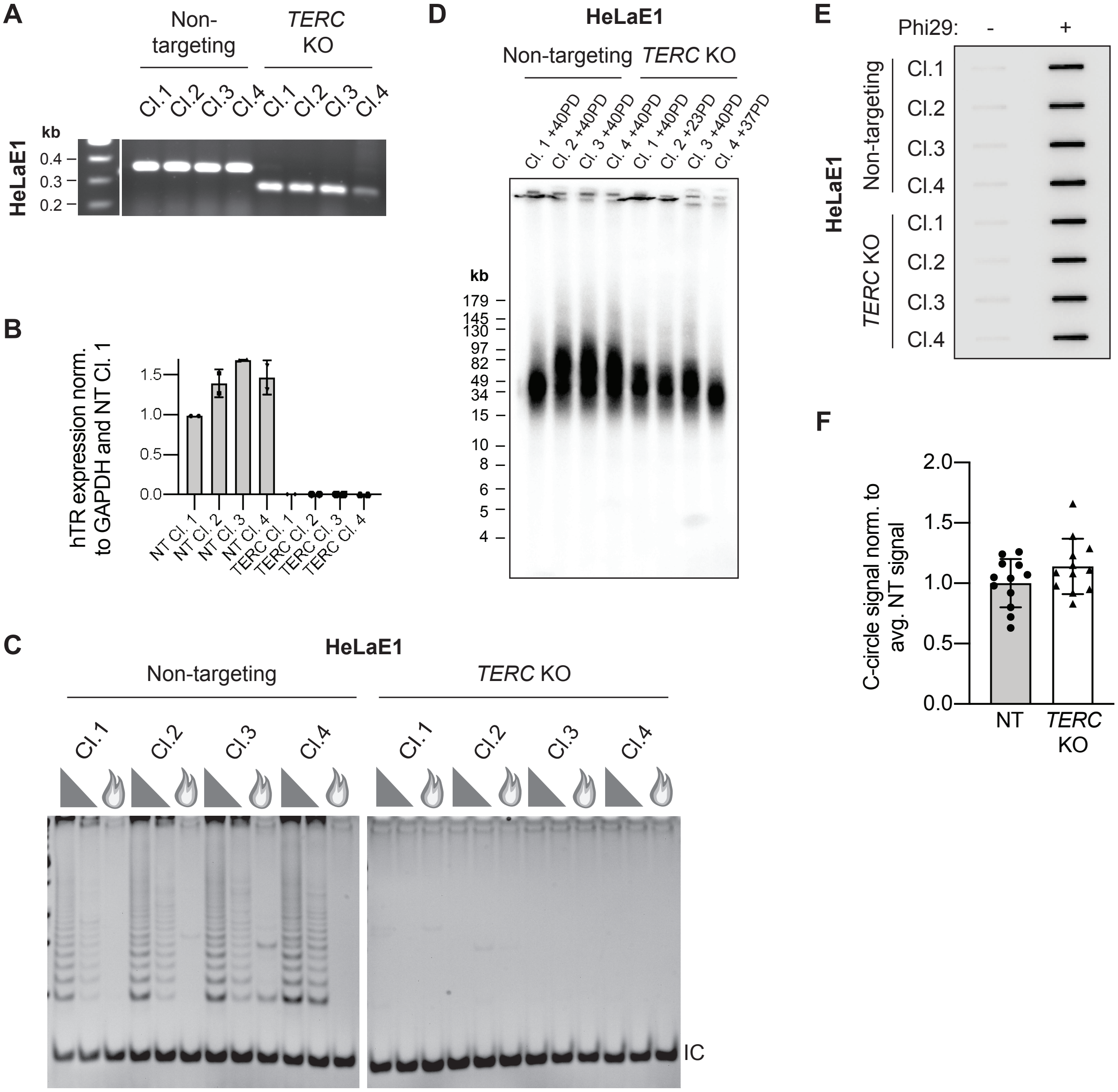


**Figure S7. *TERC* gene deletion ablates hTR expression and telomerase activity but does not impact C-circle levels in HeLa E1 cells propagated up to 40 PDs** (A) PCR amplification of the *TERC* genomic locus to assess the 84 bp deletion. The 337 bp band reflects WT sequence, and the 253 bp band reflects *TERC* KO. (B) RT-qPCR assessment of hTR expression in non-targeting (NT) and *TERC* KO HeLa E1 clones. Levels were normalized to GAPDH expression and NT clone 1. (C) TRAP assays of HeLaE1 non-targeted or TERC knockout clones. Protein concentrations used in assays as in Figure 3A. (D) TRF analysis of the non-targeting and *TERC* KO subclones as in Figure 1. The analyses were performed at PDs post-clonal isolation that were similar to those of cells collected for CCAs. The differences in telomere length in the individual clones reflects the heterogeneity of telomere lengths in the parental polyclonal populations (Figure 4A). (E) CCAs of the same clonal populations. (F) Quantification of CCAs. The signal from each clone was normalized to the average signal of the non-targeting clones per replicate. For each, four samples assayed over three experiments. Average +/- 1 standard deviation (SD). Statistical analysis was performed using an unpaired Student's t-test.
